# *Acanthamoeba polyphaga de novo* transcriptome and its dynamics during Mimivirus infection

**DOI:** 10.1101/2022.07.20.500700

**Authors:** Reut Nuri, Ester Feldmesser, Yael Fridmann-Sirkis, Hadas Keren-Shaul, Reinat Nevo, Abraham Minsky, Ziv Reich

## Abstract

*Acanthamoeba polyphaga mimivirus* (Mimivirus) is a giant virus that infects *Acanthamoeba* species – opportunistic human pathogens. We applied long- and short-read sequencing to generate a *de novo* transcriptome of the host and followed the dynamics *of both host and virus transcriptomes* over the course of infection. The assembled transcriptome of the host included 22,604 transcripts and 13,043 genes, with N50 = 2,372 nucleotides. Functional enrichment analysis revealed major changes in the host transcriptome, namely, enrichment in downregulated genes associated with cytoskeleton homeostasis and DNA replication, repair, and nucleotide synthesis. These modulations, together with those implicated by other enriched processes, indicate cell cycle arrest, an event we demonstrated experimentally. We also observed upregulation of host genes associated with transcription, secretory pathways and, as reported here for the first time, peroxisomes and the ubiquitin-proteasome system. In Mimivirus, the early stages of infections were marked by upregulated genes related to DNA replication, transcription, translation, and nucleotide metabolism, and the later stages, by enrichment in genes associated with lipids metabolism, carbohydrates, and proteases. Some of the changes observed in the amoebal transcriptome likely point to Mimivirus infection causing the dismantling of the host cytoskeleton, the translocation of endoplasmic reticulum membranes to viral factory areas, and cell cycle arrest.

## Introduction

*Acanthaomeba polyphaga mimivirus* (Mimivirus) is a member of the nucleo-cytoplasmatic large DNA viruses (NCLDV) clade, whose infection of *Acanthamoeba spp*. causes massive lysis in the amoebal population. The first giant DNA virus to be discovered, Mimivirus has a diameter of approximately 750 nm. Its genome, ∼1.2 × 10^6^ base-pair (bp) long, encodes approximately 1000 open reading frames (ORFs), many of which are orphan genes, with no detectable homologues in other species [1–4].

Mimivirus’s target hosts*, Acanthamoeba* spp., are free-living amoebae that reside in air, soil, and water environments and are recognized as opportunistic human pathogens [5–7]. Indeed, *Acanthaomeba* polyphaga (AP) is a causative agent of several human diseases, including granulomatous amoebic encephalitis, a rare brain infection that is generally fatal, a skin ulcer in human immunodeficiency virus (HIV) patients, and Acanthamoeba keratitis - a corneal infection that can lead to loss of vision [6]. The latter is becoming more prevalent because of the increased use of contact lenses, which nurture amoebal growth when hygiene is inadequate.

Mimivirus enters amoeba host cell via phagocytosis. In the phagosome, it loses its capsid and fuses with the phagosome membrane, after which the viral core enters the cytoplasm [8, 9]. Therein, viral factories (VFs) begin to be formed and, at 5 hours post infection (HPI), they fuse into a single mature VF [8, 10, 11]. Viral production occurs in the VF and, at approximately 12–14 HPI, cell lysis occurs, prompting the release of newly-formed viruses into the surrounding medium [8, 11].

To date, one work has described Mimivirus transcriptome dynamics during the infection cycle in *Acanthamoeba castellanii* (AC), another member of the *Acanthamoeba* genus [12]. However, the dynamics of the host transcriptome over the course of infection has not yet been characterized. In this study, the AP transcriptome was followed throughout the different phases of infection. Since no transcriptome was available for AP, we assembled its transcriptome *de novo* using PacBio long-read (LR) sequencing and Illumina sequencing of cDNA produced from RNA from uninfected AP and infected AP 1, 3, and 5 HPI, corresponding to early, intermediate, and the beginning of the late stage of infection, respectively. A functional enrichment analysis of AP and Mimivirus gene expression was performed at the different time points. The analysis highlights biological processes and organelles in AP whose related genes are enriched during the infection cycle, including peroxisome lipid transport, peroxisomal organization and the ubiquitin-proteasome system. Other enriched genes can be linked to the dismantling of the cytoskeleton [11], translocation of ER membranes to the periphery of the VF [13, 14], and cell-cycle arrest. The latter was observed experimentally. We also demonstrate here, *de novo* nucleotide synthesis during Mimivirus infection, despite enrichment in host genes that downregulate nucleotide synthesis.

## Results

### De novo assembly and processing of the Acanthamoeba polyphaga transcriptome

#### Overview of the transcriptomics analysis process

To obtain the AP transcriptome *de novo*, we performed PacBio Single Molecule Real-Time (SMRT) sequencing for long reads (LRs) and Illumina next-generation sequencing (NGS) of AP samples after 1, 3, and 5 hours of infection with Mimivirus, as well as of uninfected AP cells and AP cells 1 h after UV treatment. The latter analysis served to compare the stress induced by viral infection to abiotic stresses.

For transcriptome assembly, 147 million Illumina paired short reads (SRs) and 116,781 circular consensus sequences from PacBio sequencing were used. LR-based transcripts were used as the basis of the transcriptome. SRs were mapped to the LRs, and unmapped reads were assembled using genome-based and non-reference-based *de novo* assembly tools. The assembly resulted in 22,604 transcripts and 13,043 genes.

Benchmarking Universal Single-Copy Ortholog (BUSCO) scores were used to assess the completeness of the transcriptome. Selected transcripts were Sanger-sequenced to verify accuracy. Host differential expression analysis was performed by comparing gene expression levels at each time point with their levels in the uninfected sample. Overall, 5,344 differentially expressed genes (DEGs) (40.9%) were detected in the host during the infection. The transcripts were annotated using the Gene Ontology (GO) annotation system [15], which was used for the functional enrichment analysis. Mimivirus differential expression analysis was performed between any two of the three time points (1 HPI vs. 3 HPI, 1 HPI vs. 5 HPI and 3 HPI vs. 5 HPI) and yielded 870 DEGs (90.8%), which were assigned to three clusters using k-mean clustering. These clusters were later used for enrichment analysis based on manual gene categorization. A detailed description of these steps and their results follows.

### AP transcriptome assembly and processing

PacBio LRs, processed using IsoSeq3 [16], were used as the basis for the construction of the *de novo* transcriptome. Paired reads from Illumina sequencing were then processed and subsequently mapped to the LRs using Bowtie2 [17, 18]. Unmapped reads were used to build transcripts that were not covered by the SMRT sequencing. Two assembly algorithms were used: i) StringTie [19, 20], based on the AP genome; and ii) rnaSPAdes [21, 22], for non-database-based assembly. Transcripts derived from the three sources (Isoseq3, StringTie, and rnaSPAdes) were combined into a single unified transcriptome. The united transcriptome was subjected to redundancy removal, based on the longest coding sequence, using EvidentialGene [23, 24] (Figure 1). Blastx [25] was used to remove Mimivirus transcripts from the assembled transcriptome, as well as to remove contaminations by other organisms that may reside in the amoeba [26] or may have been introduced during sample preparation.

**Figure 1.**
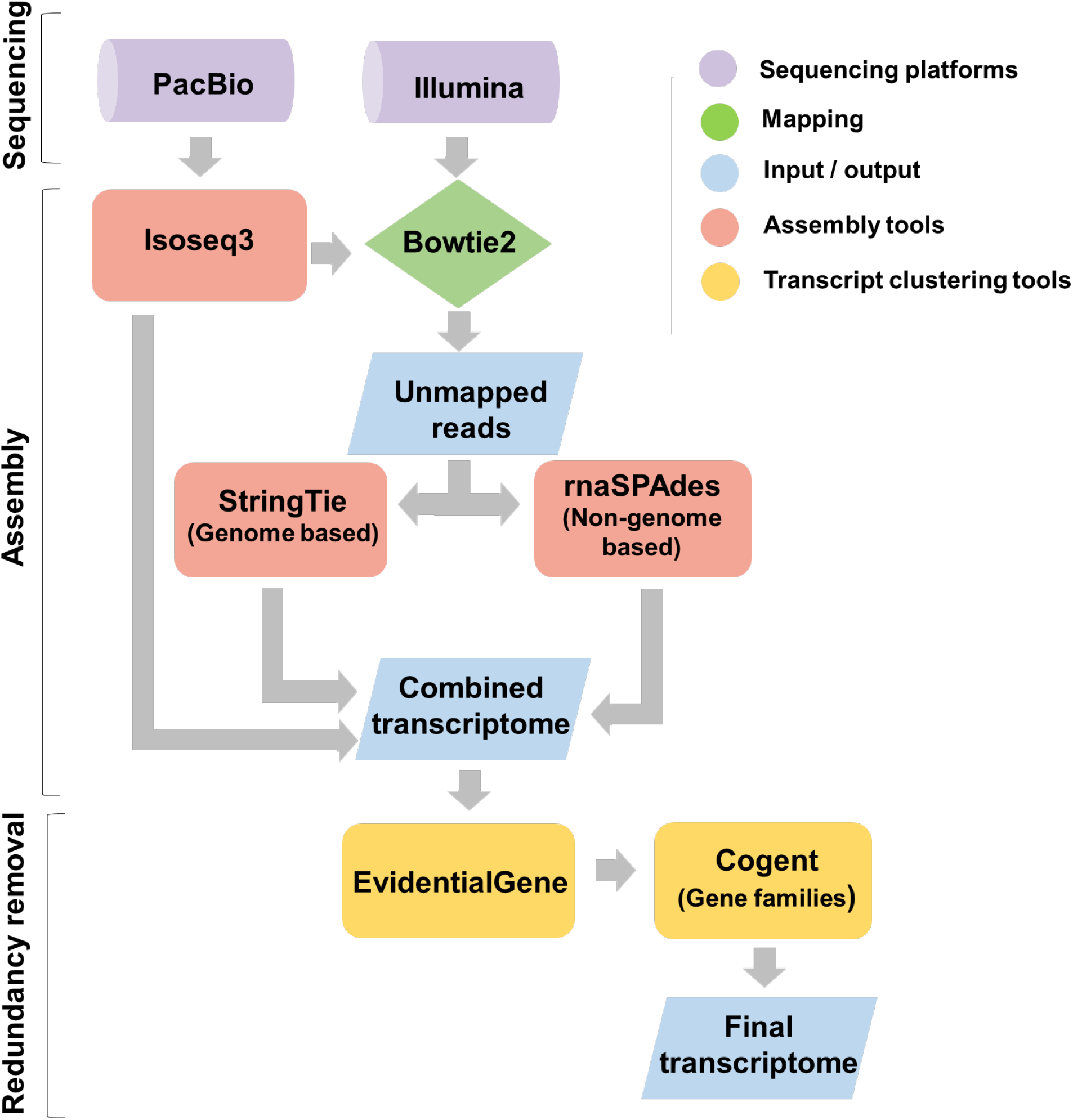
Transcriptome assembly workflow. Illumina NGS was used for short-read (SR) cDNA sequencing, and PacBio SMRT for long-read (LR) cDNA sequencing. PacBio results were processed using the Isoseq3 tool. SRs were mapped to LRs using the Bowtie2 tool [69]. Unmapped reads were used to further assemble the transcriptome via StringTie [19, 20] (genome-based) and rnaSPAdes [21, 22]. Transcripts from Isoseq3, StringTie and rnaSPAdes were combined, and all the transcripts were processed using EvidentialGene [15], to remove redundancy, and the Cogent tool [19], to create gene families out of isoforms.

#### Transcriptome characteristics

The assembled AP transcriptome yielded 22,604 transcripts (Supplementary File 1), with a length distribution as indicated in Figure 2A and N50 = 2,372 bp. Collapsing the transcripts of the same gene to the corresponding gene family (Supplementary Table 1), using Cogent [27], resulted in 13,043 genes, most of which had a single isoform (Figure 2B). In addition, we used the transcriptome as the basis for the AP proteome, which we also show here (Supplementary File 2). The species for which matching annotations were obtained were analyzed based on the top Blastx results (Supplementary Table 2). As expected [28], approximately 82% of the genes with taxonomy assignment matched to the *Acanthamoeba castellanii* transcriptome (Figure 2C), a companion species in the *Acanthamoeba* genus. All top-five organisms from which annotations were borrowed were eukaryotes; annotations of 394 genes (4%) originated from bacteria and 15 genes (0.15%) from archaea. Assigning genes to GO terms using Blast2go [29] yielded 9,921 genes with GO annotations (76% of the genes), which were then used for the enrichment analysis (Supplementary File 3).

**Figure 2.**
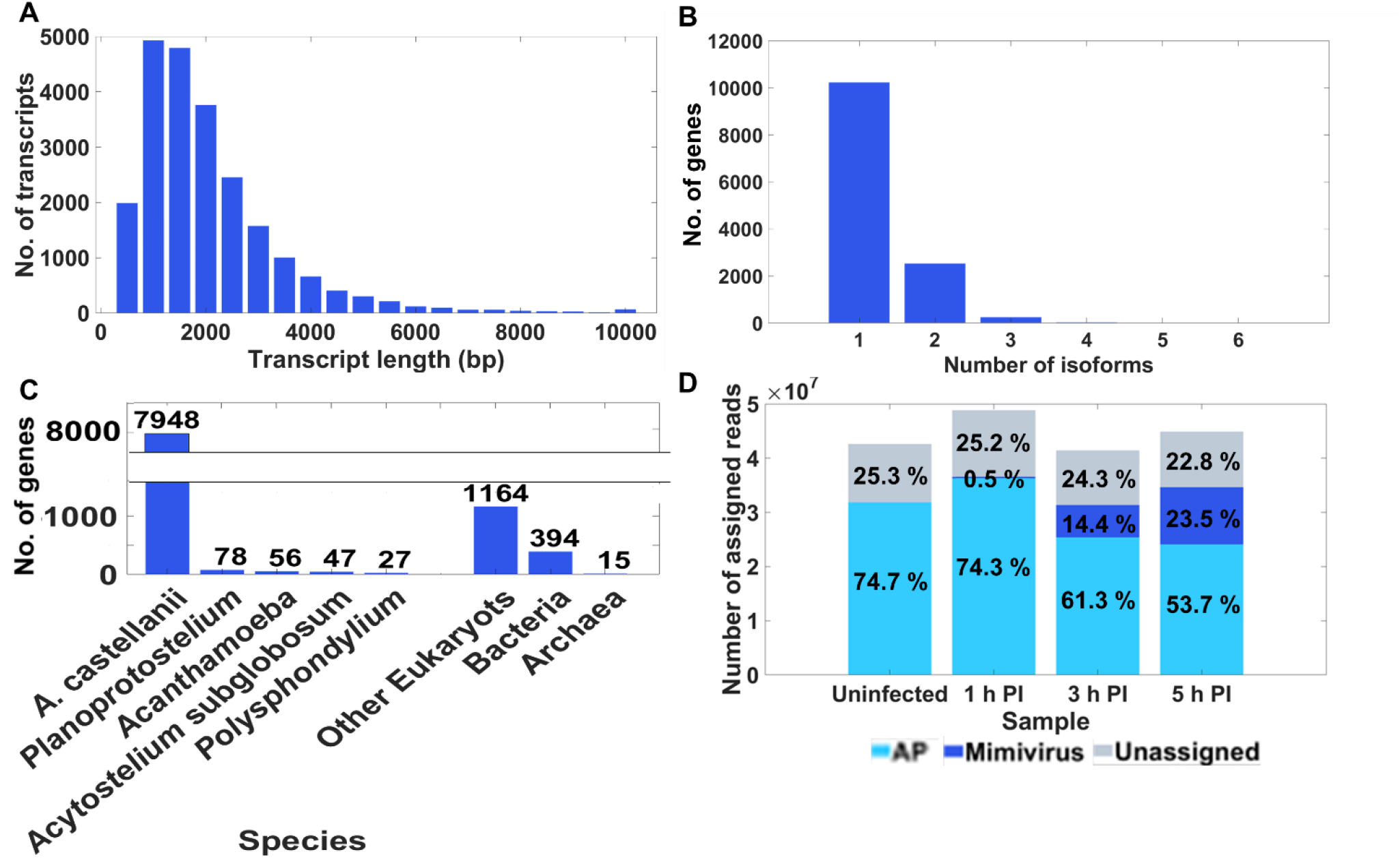
Transcriptome characteristics. A. Distribution of transcript size (nucleotides). B. Distribution of the number of isoforms per gene. C. Taxonomic allocation of borrowed annotations. The top 5 organisms from which gene annotations were borrowed are shown in the left five bars. The rest of the genes were classified as eukaryotes, prokaryotes or archaea (three right bars). D. Read distribution between *Acanthaomeba polyphaga* (AP) and *Acanthaomeba polyphaga mimivirus* (Mimivirus) in the different samples. The percentage from total reads at a certain time point is indicated on the bars.

As infection progressed, the percentage of Mimivirus-assigned reads in the ensemble rose from 0.5 at 1 HPI, to 14.4 at 3 HPI and 23.5 at 5 HPI (Figure 2D), while the percentage of AP assigned reads declined.

Transcription factor (TF) analysis was performed using HMMER [30], running on animal and plant TF databases. The analysis yielded 280 candidate TFs in the AP transcriptome (Supplementary Table 3), exhibiting various expression levels at different time points along the infection. For comparison, the number of TFs identified in other classes of the Amoebozoa phylum, *Entamoeba histolytica* and *Dictyostelium* spp., are 69, 116 and 123, respectively.

To obtain information about splice junction sequences and intron lengths, transcripts were mapped to the available AP genome [31], using Minimap2 [32, 33]. AP appears to have a canonical splice junction sequence identical to that of other eukaryotes, such as fungi, plants and animals [34], with GT and AG dinucleotides at the donor and acceptor splice sites, respectively (Figure 3A). The median length of AP introns is 72 bp (Figure 3B), similar to that in *Entamoeba histolytica* (75 bp) [35].

**Figure 3.**
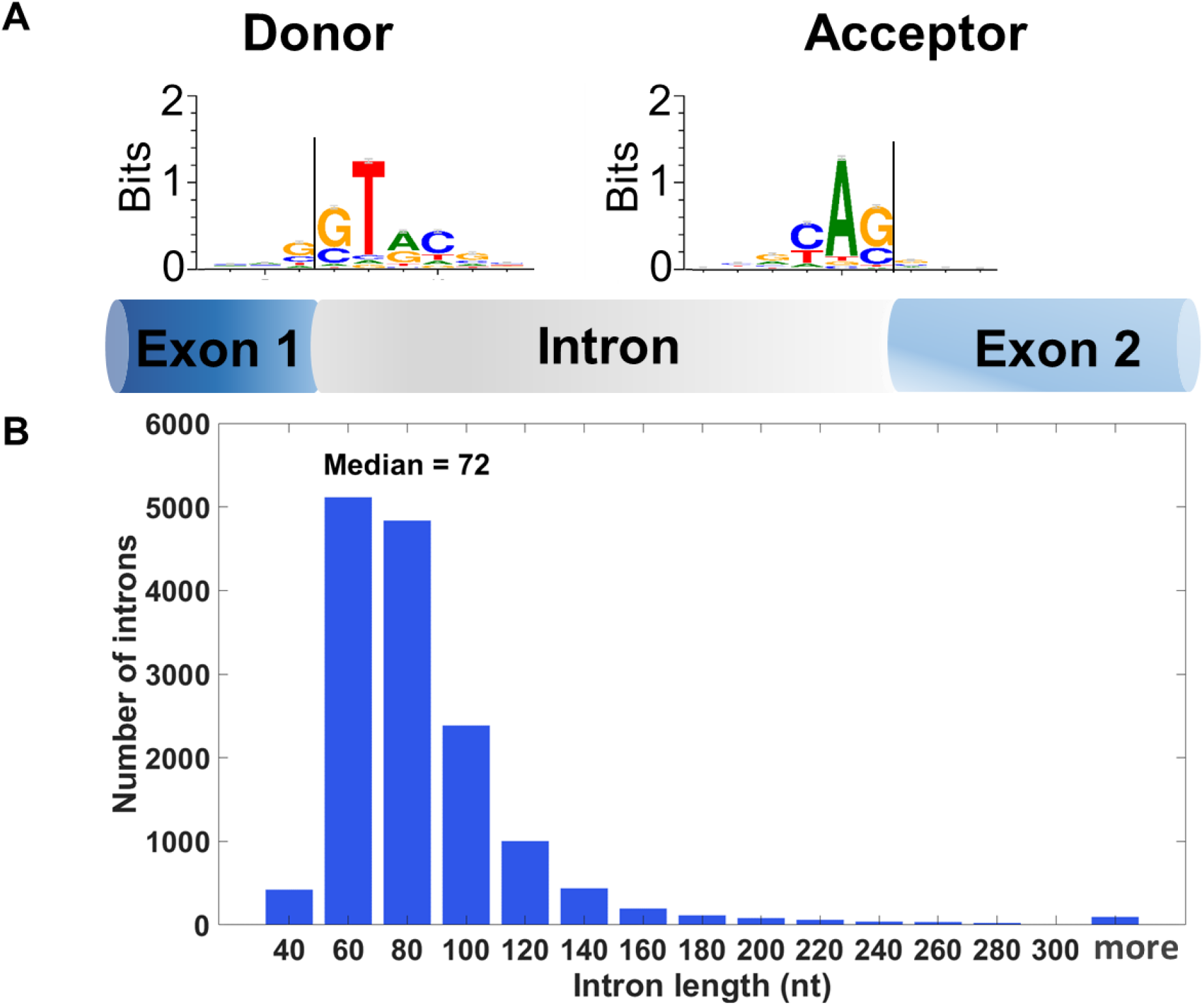
Splice junction analysis of *Acanthaomeba polyphaga* (AP). A. Consensus sequence of donor and acceptor splice sites of AP, as determined from long read transcripts, presented graphically in the form of a logo consisting of stacks of single-letter nucleic acid symbols. The larger the letter in each stack, the greater the frequency of the corresponding nucleic acid at that position in the sequence. B. Distribution of intron lengths (in base pairs).

#### Quality control of transcriptome assembly and accuracy

Principal component analysis (PCA) of both AP and Mimivirus gene expression levels was performed to compare the biological duplicates and the samples collected from the different time points along the infection (Supplementary Figure 1). The results indicated that the duplicates of each sample were similar. The uninfected and 1 HPI samples were also similar, whereas there were significant differences in gene expression along the infection process between samples obtained at the other time points.

In addition, we used BUSCO [36, 37] to assess the completeness of the transcriptome. This tool uses a list of genes that are common to a certain taxon and are present in a single copy and searches for them in the database. Since BUSCO is a robust comparative tool, we compared BUSCO scores derived from the assembled AP transcriptome with those obtained from the transcriptome of AC [38]. Results obtained utilizing the eukaryotic BUSCO database showed the AP transcriptome to be more complete and with more duplications than the AC transcriptome (Table 1). Running the two transcriptomes against the protist BUSCO-database produced a similar trend (Supplementary Table 4). Since the Mimivirus transcriptome is known, we used it as an independent means to assess the quality of the assembly, and Mimivirus-originated transcripts were aligned to the published Mimivirus transcriptome. Alignment resulted in 86.2% completeness, with 15.5% duplications (Supplementary Table 5). To examine the accuracy of the transcripts, we performed sample testing by Sanger sequencing of 13 randomly selected transcripts from the different assemblers (Isoseq3, StringTie and rnaSPAdes). This analysis included 99.1–100% identity with transcript sequences for 11 of the transcripts. The coverage alignments were 55.6 and 90% for two of the four transcripts that originated from rnaSPAdes, a non-database-based assembly (Supplementary Table 6).

**Table 1:**
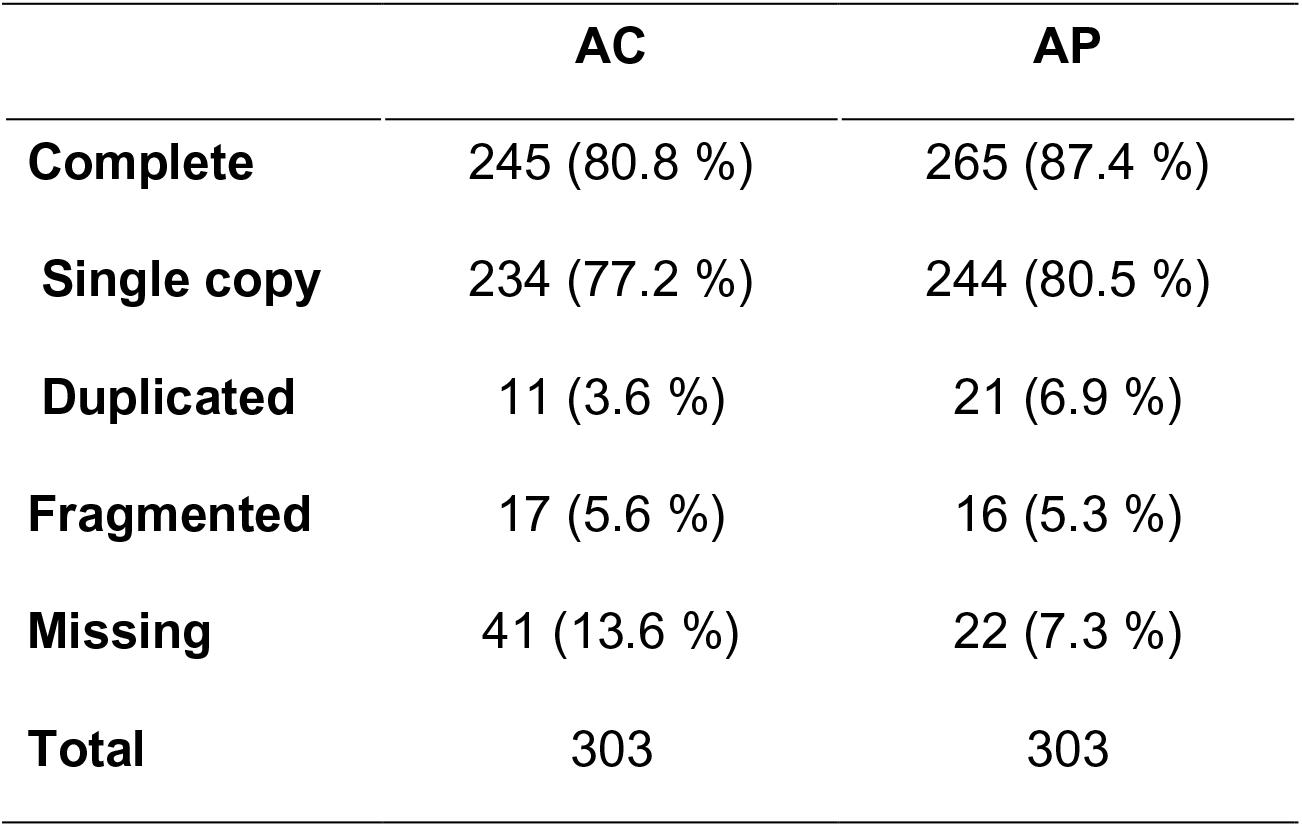
BUSCO of AC and AP transcriptomes against the BUSCO eukaryotic database

|  | **AC** | **AP** |
| --- | --- | --- |
| **Complete** | 245 (80.8 %) | 265 (87.4 %) |
| **Single copy** | 234 (77.2 %) | 244 (80.5 %) |
| **Duplicated** | 11 (3.6 %) | 21 (6.9 %) |
| **Fragmented** | 17 (5.6 %) | 16 (5.3 %) |
| **Missing** | 41 (13.6 %) | 22 (7.3 %) |
| **Total** | 303 | 303 |

### Functional enrichment analysis of the Mimivirus transcriptome

The Mimivirus DEGs (Supplementary Table 7) are divided into three clusters. (Supplementary Table 8). These clusters represent the expression of genes in early (208 DEGs), intermediate (274 DEGs) and late (388 DEGs) stages of infection (Figure 4). Among the detected genes, 115 maintained approximately constant expression levels throughout the examined time points.

**Figure 4.**
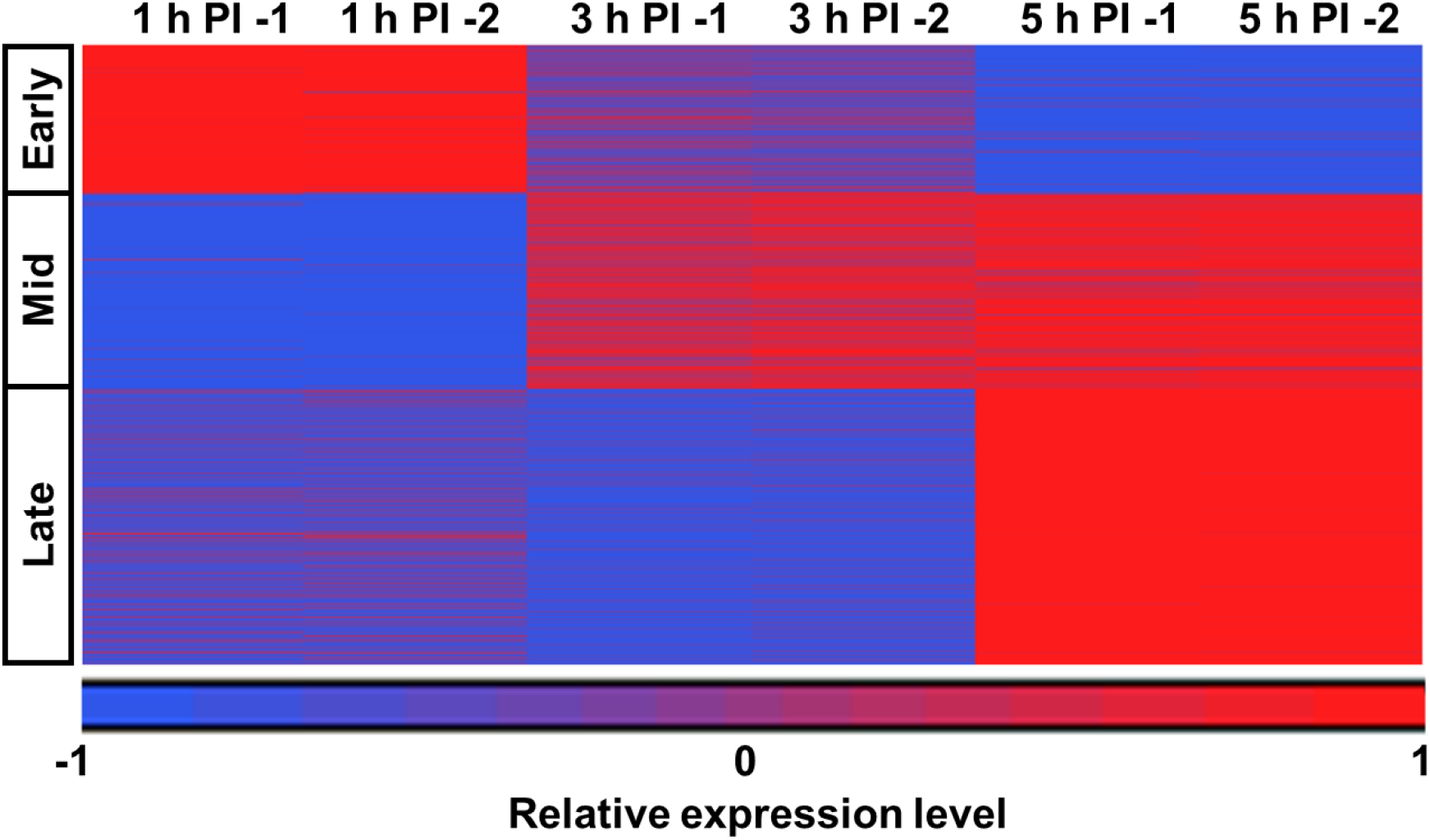
K-mean clustering of the differentially expressed genes (DEGs) of *Acanthaomeba polyphaga mimivirus* (Mimivirus). Mimivirus DEGs were classified into three clusters using k-mean clustering, as was suggested by the Silhouette score (***Supplementary Figure 2***). These clusters correspond to genes expressed in the early, intermediate, and late infection stages.

Since only a relatively small number of Mimivirus genes have GO annotations, we manually categorized the genes according to the top Blastx results and the literature (Supplementary Table 8). A statistical analysis of the data using a hypergeometric test indicated enriched categories in each examined time point (Figure 5A) (Supplementary Table 9).

**Figure 5.**
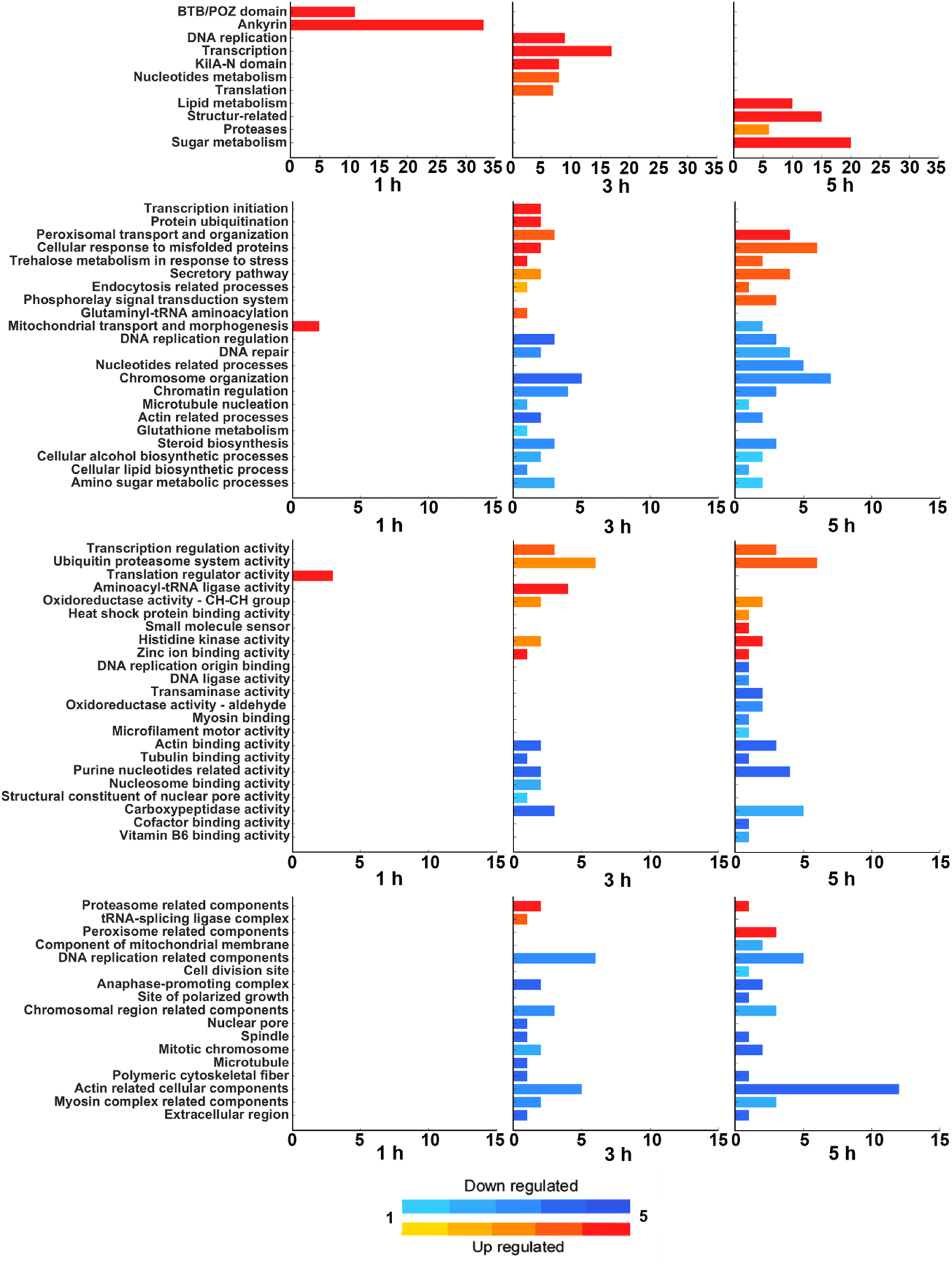
Functional enrichment analysis. Functional enrichment analysis of *Acanthaomeba polyphaga mimivirus* (Mimivirus) (A) and *Acanthaomeba polyphaga* (AP) (B–D). For the simpler Mimivirus, each category comprises a group of genes, with the bar size proportional to the number of genes in the category. For AP, the genes are annotated to each GO term, with groups of GO terms subsequently semi-automatically grouped into categories. Each category is represented by a bar whose size is proportional to the number of GO terms it includes. The functional enrichment analysis is presented with respect to the three principal GO domains: (B) biological processes; (C) molecular function; and (D) cellular components. The coloring of the bars indicates whether the category became enriched in up-regulated (red) or down-regulated (blue) genes over the course of infection. The hue represents the significance of the enrichment; in A, it is the p-value of the enriched category, and in B-D, it is based on the average p-value of the GO terms in the category.

The first cluster contained genes with the highest expression level at 1 HPI. Here, enrichment was noted for the Mimivirus ankyrin genes and TB/POZ-domain-containing proteins (Figure 5A). The ankyrins represent the largest category in the Mimivirus transcriptome, consisting of 106 putative genes. Of these, 33 were found in the 1 HPI cluster. Ankyrin repeat-containing proteins are known to be involved in protein-protein interactions [39], though no specific function of ankyrins has been demonstrated in Mimivirus.

The genes in the second cluster were highly expressed at 3 HPI and most were also highly expressed at 5 HPI (Figure 4). Specifically, Mimivirus genes for KilA-N domain, DNA replication, transcription, nucleotide metabolism and translation were enriched (Figure 5A), along with the corresponding AP categories (Figure 5B-D).

The third cluster was dominated by genes that had the highest expression level at 5 HPI. These included genes encoding for proteases, structural proteins, and lipid- and sugar-metabolism-related proteins (Figure 5A). The lipid metabolism category consists of genes encoding for proteins putatively involved in lipid modification, binding, transport, and processing; the structure-related category includes, among other genes, eight putative collagen genes (collagen 1–8) present in the Mimivirus genome (L71, R196, R239, R240, R241, L668, L669 and L623).

It was reported that Mimivirus undergoes large-scale glycosylation on its fibrils [40, 41] and possesses various genes that encode for proteins involved in glycosylation [12, 41–43]. In line, twenty of the 22 genes classified to the sugar metabolism category were among those assigned to the 5 HPI cluster. Many of these genes are related to glycosylation (L619, L373, R707, L140, L263, R363, L137, L138 and L193).

### Functional enrichment analysis of the AP transcriptome

To gain insight into the dynamics of the AP transcriptome during the different Mimivirus infection stages, we selected genes that were differentially expressed at each of the time points compared to their expression in the uninfected ameba (Supplementary Table 10). We then divided the DEGs into up-regulated (UR) and down-regulated (DR) genes for each time point, which yielded six gene groups (1 h UR or DR, 3 h UR or DR and 5 h UR or DR). The different groups were analyzed using Ontologizer [44, 45] (Figure 5B-D). We used the three primary domains of the GO acyclic tree: biological processes (Figure 5B), molecular function (Figure 5C) and cellular components (Figure 5D). The GO terms belonging to each domain were grouped into categories in a semi-automatic manner (Supplementary Table 11).

Only 31 DEGs were identified at 1 HPI compared to the uninfected sample. The high similarity between the two samples can also be inferred from the PCA analysis (Supplementary Figure 1). By comparison, 3,993 DEGs were identified at 3 HPI, and 4,025 DEGs at 5 HPI. Since most of the enriched categories were common to both 3 and 5 HPI (Figure 5A-C), in the subsequent sections, we discuss GO terms related to these categories together. A full list of GO terms assigned to categories and time points can be found in Supplementary Table 11.

#### Mitochondrial activity

The mitochondrial transport and morphogenesis category was enriched at 1 HPI. UR genes annotated to the respiratory chain complex III assembly (GO:0017062) and mitochondrial respiratory chain complex assembly (GO:0033108) GO terms were enriched at this time point. At 5 HPI, this category was enriched in DR genes, specifically with mitochondrial transmembrane transport (GO:1990542) and mitochondrion morphogenesis (GO:0070584) GO terms. These observations corroborate the finding of Legendre et al. [12] of an initial increase and then, later on in the infection, a decrease in the number of reads that originate from amoebal mitochondria-related genes.

#### Translation regulation

Enrichment in UR genes was observed at 1 HPI for the translation regulator activity nucleic acid binding (GO:0090079), translation factor activity RNA binding (GO:0008135), and translation regulator activity (GO:0045182) GO terms. Subsequently, at 3 and 5 HPI, the categories enriched in UR genes were translation-related glutaminyl-tRNA aminoacylation (GO:0006425) (Figure 5B) and aminoacyl-tRNA ligase activity (Figure 5C). The tRNA-splicing ligase complex category (Figure 5D) was enriched in UR genes at 3 HPI but not at 5 HPI. Unlike most viruses, Mimivirus encodes for translation-related proteins in its genome. The Mimivirus translation-related category was enriched at 3 HPI (Figure 5A) with genes encoding for translation initiation factors (R458, L529, L496), tRNA ligases (L164, L124) and methionyl-tRNA synthetase (R639).

#### Transcription

At 3 and 5 HPI, the AP transcriptome was enriched in UR genes associated with transcription initiation, including terms related to the regulation of DNA-templated transcription initiation (GO:2000142) and regulation of transcription initiation from RNA polymerase II promoter (GO:0060260) (Figure 5B). This was accompanied by the UR of transcription regulation activity (Figure 5C), including DNA-binding transcription factor activity (GO:0003700) and general transcription initiation factor binding (GO:0140296) GO terms.

In the Mimivirus, the transcription category was also enriched at 3 HPI (Figure 5A), consistent with a previous report of transcription activity in VFs at 4 HPI [46]. Among the genes belonging to this category were transcription regulators L544, R450, R453, R430 and R339; DNA-directed RNA polymerases R501, R867, L244, L235, R209, R470, L208, R357b and L376; the two mRNA capping enzyme-encoding genes R382 and L308; and the poly-A polymerase catalytic subunit L341 (Note that some of the Mimivirus transcription-related proteins are also packed in the virion [10] and might play a role during earlier infection stages). Another pertinent Mimivirus-enriched category was proteins containing KilA N (Figure 5A), a DNA-binding domain found in bacteria and DNA viruses [47].

### Peroxisomes

Found in most eukaryotic cells, peroxisomes are small organelles that carry out various oxidative-related reactions. A significant function of peroxisomes is related to lipid metabolism, including the breakdown of fatty acids and the synthesis of very-long chain fatty acids and ether lipids [48]. Our data reveal enrichment in UR genes related to peroxisomal transport and organization (Figure 5B) at both 3 and 5 HPI. Go terms belonging to this category include long-chain fatty acid import into peroxisome (GO:0015910) and peroxisomal membrane transport (GO:0015919). Terms associated with peroxisome-related components (Figure 5D) were enriched in UR genes solely at 5 HPI. These GO terms include the integral component of peroxisomal membrane (GO:0005779) and the peroxisome (GO:0005777).

### The secretory pathway and the ubiquitin-proteasome system

The ER is a major component of the secretory pathway. Its membrane-bound lamellar network, which is continuous with the outer nuclear envelop, constitutes more than half the mass of the cell’s membrane [48]. Its primary roles are post-translational protein modification and lipid synthesis. Host GO terms related to the secretory pathway were enriched in UR genes at 3 and 5 HPI (Figure 5B). These included ER-to-cytosol transport (GO:1903513) and the vesicle targeting and rough ER to cis-Golgi (GO:0048207) GO terms.

The proteasome is a large protein complex that degrades ubiquitin-tagged proteins. This proteolytic activity plays vital roles in various cellular processes such as transcription, cell cycle progression and cell homeostasis [49]. At 3 HPI, there was enrichment in UR genes related to the protein ubiquitination category (Figure 5B), which includes protein modification by small protein conjugation (GO:0032446) and protein ubiquitination (GO:0016567) GO terms. In addition, at both 3 and 5 HPI, there was enrichment in UR genes associated with the ubiquitin-proteasome system activity category (Figure 5C), including poly-ubiquitin modification-dependent protein binding (GO:0031593), proteasome binding (GO:0070628) and ubiquitin-protein transferase activity (GO:0004842) GO terms. Also enriched in UR genes were GO terms associated with the proteasome-related components category (Figure 5D), including the proteasome accessory complex (GO:0022624) and the proteasome regulatory particle (GO:0005838). Another category enriched in UR genes at 3 and 5 HPI that is related to both the ubiquitin-proteasome system and ER is the cellular response to misfolded proteins category (Figure 5B). This category includes cellular response to heat (GO:0034605), response to topologically incorrect protein (GO:0035966), and ubiquitin-dependent ERAD pathway (GO:0030433) GO terms.

Interestingly, the Mimivirus itself appears to possess genes encoding for components of the ubiquitin-proteasome system, such as putative E2/E3 enzymes (Supplementary Table 8). This Mimivirus gene category, however, was expressed uniformly throughout the infection cycle and was not enriched at any of the examined time points.

### Cell Cycle

During Mimivirus infection, AP cells stop dividing (Supplementary Video 1). This is also evidenced by the transcriptomic results, as many GO terms related to cell cycle categories were enriched in DRGs starting at 3 HPI. These terms include DNA replication origin binding, DNA repair, chromatin regulation, chromosome organization, DNA replication-related components, and anaphase-promoting complex (Figure 5C, D). A related category, DNA replication origin binding activity (Figure 5B), was enriched in DRGs only at 5 HPI. Some DRG GO terms associated with the two categories are shown in Table 2 (A full list of associated GO terms is provided in Supplementary Table 2).

**Table 2:** Host DNA replication-related categories and examples of associated GO terms that exhibit enrichment in DR genes at 3 and 5 HPI

| AP category enriched in DR genes | Associated GO terms (examples) |  |
| --- | --- | --- |
|  | Name | Number |
| **At both 3 and 5 HPI** |  |  |
| DNA-replication regulation | DNA replication-independent nucleosome organization | GO:0034724 |
|  | DNA strand elongation | GO:0022616 |
|  | DNA-replication initiation | GO:0006270 |
| DNA-replication-related components | MCM complex | GO:0042555 |
|  | Ctf18 RFC-like complex | GO:0031390 |
|  | DNA polymerase complex | GO:0042575 |
|  | DNA-replication preinitiation complex | GO:0031261 |
| DNA repair | Double-strand break repair via break-induced replication | GO:0000727 |
|  | DNA ligation involved in DNA repair | GO:0051103 |
|  | Negative regulation of DNA recombination | GO:0045910 |
| Chromatin regulation | Chromatin silencing | GO:0006342 |
|  | Chromosome condensation | GO:0030261 |
|  | Nucleosome assembly | GO:0006334 |
| Chromosome organization | Anaphase-promoting complex-dependent catabolic process | GO:0031145 |
|  | Sister chromatid cohesion | GO:0007062 |
|  | Mitotic sister chromatid separation | GO:0051306 |
| Anaphase-promoting complex | Anaphase-promoting complex | GO:0005680 |
|  | Nuclear ubiquitin ligase complex | GO:0000152 |
| Purine-nucleotide-related activity | Guanyl-nucleotide exchange factor activity | GO:0005085 |
|  | Carboxypeptidase activity | GO:0004180 |
|  | Adenine nucleotide transmembrane transporter activity | GO:0000295 |
|  | Purine ribonucleotide transmembrane transporter activity | GO:0005346 |
| **At 5 HPI only** |  |  |
| DNA replication origin binding | DNA replication origin binding | GO:0003688 |
| Nucleotide-related processes | Adenine nucleotide transport | GO:0051503 |
|  | Purine ribonucleotide transport | GO:0015868 |

In marked contrast, the viral transcriptome exhibited enrichment in UR DNA replication-related genes at 3 HPI (Figure 5A). Amongst these were mostly genes encoding for DNA polymerases (R322, R493) and replication factors (R411, L478, L499, R395 and R510). Other UR genes associated with DNA replication were enriched at 5 HPI. These included primase polymerase (L537) and a member of DNA polymerase family X (L318). Interestingly, replication of Mimivirus DNA was observed already at 2 HPI [46]. This early phase is likely mediated by DNA replication-related proteins, e.g., R303, L478 R493, R771 and others, that are packed in the virion [3, 10] and presumably enable the commencement of replication [12].

Other host categories enriched in DRGs were nucleotide-related processes, at 5 HPI (Figure 5B), and purine nucleotide-related activity, at both 3 and 5 HPI (Figure 5C) (Table 2). Unlike AP, the Mimivirus category of nucleotide metabolism was enriched at 3 HPI (Figure 5A). To identify the source of nucleotides used for the replication of the viral genome, the host cells were treated with a nucleoside analog of thymidine prior to infection. The analog, 5- ethynyl-2’-deoxyuridine (EdU), can be fluorescently labeled after fixation and permeabilization of the cells. EdU was incorporated only into the host genome, thus enabling to distinguish between existing and *de novo-*synthetized nucleotides. While we observed prominent staining by the DNA dye 4′,6- diamidino-2-phenylindole (DAPI) in the VFs following infection, it was not accompanied by EdU staining (Figure 6). Thus, nucleotides used during Mimivirus genome replication originated primarily or solely from *de novo* synthesis, where viral proteins are likely to play a major role.

**Figure 6.**
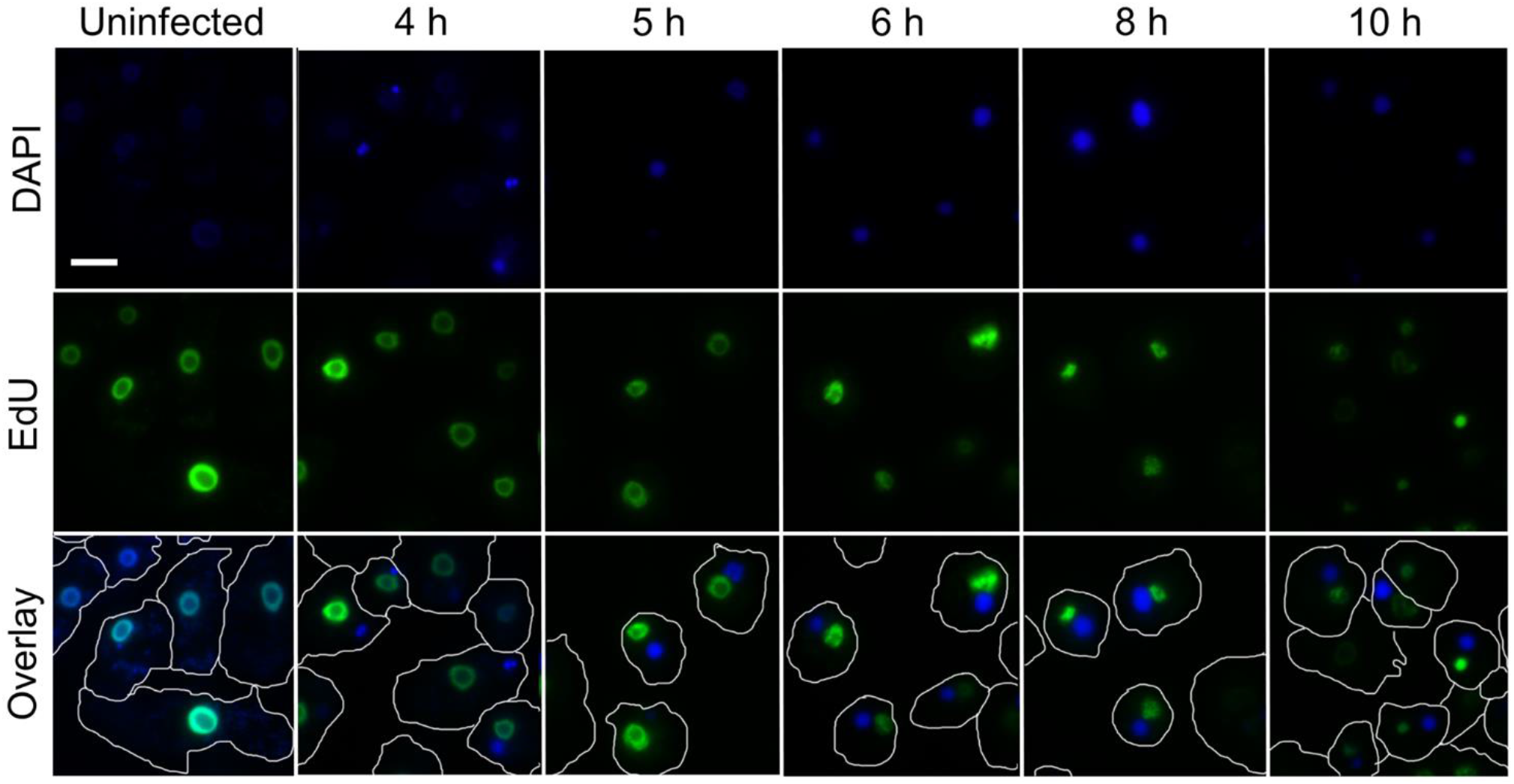
Nuclear DNA staining with EDU-labeled nucleotides. AP cells were grown in PYG medium supplemented with 20 µM EDU-labeled nucleotides for approximately 24 h prior to infection. One hour before the infection, the medium was replaced with fresh PYG. The AP cells were infected for the indicated time periods, after which cells were fixed and stained. Green - EDU labeling (nuclear DNA). Blue - DAPI staining. White line - cell contour. Scale bar: 10µm.

### Cytoskeleton

Similarly to other eukaryotes, AP possesses an extensive cytoskeletal network that confers mechanical stability to the cells and plays important roles in cellular transport and locomotion [5]. Mimivirus infection induces the rearrangement of tubulin and actin networks within AP cells [11] and causes them to lose their asymmetrical shape with pseudopodia and begin to round, with their actin fibers retracting to form ‘shell’-like structures at the cell periphery [11]. Consistent with these observations, we found that categories and associated GO terms related to cytoskeletal components were enriched in DRGs upon infection (see Table 3 for principal examples and Supplementary Table 11 for the full list). Specifically, both at 3 and 5 HPI, there was enrichment in DRGs in the microtubule nucleation (Figure 5B), tubulin binding activity (Figure 5C), and microtubule (Figure 5D) categories. Likewise, actin-related processes (Figure 5B), actin binding (Figure 5C) and actin-related cellular components (Figure 5D) all showed enrichment in DRGs. The myosin binding (Figure 5C) and myosin complex (Figure 5D) categories were also enriched in DRGs at 3 and 5 HPI.

**Table 3:** Host cytoskeleton-related categories and examples of associated GO terms that exhibit enrichment in down regulated (DR) genes 3 and 5 HPI

| AP category enriched in DR genes | Associated GO terms (examples) |  |
| --- | --- | --- |
|  | Name | Number |
| Microtubule nucleation | Microtubule nucleation | GO:0007020 |
| Tubulin binding activity | Tubulin binding | GO:0015631 |
| Microtubule | Microtubule | GO:0005874 |
| Actin-related processes | Negative regulation of cytoskeleton organization | GO:0051494 |
|  | Actin crosslink formation | GO:0051764 |
| Actin binding | Actin filament binding | GO:0051015 |
| Actin-related cellular components | Cortical cytoskeleton | GO:0030863 |
|  | Lamellipodium | GO:0030027 |
|  | Cortical actin cytoskeleton | GO:0030864 |
| Myosin binding | Myosin binding | GO:0017022 |
| Myosin complex | Actomyosin | GO:0042641 |
|  | Myosin I complex | GO:0045160 |

## Discussion

In this work, we generated the transcriptome of the free-living amoeba *Acanthamoeba polyphaga* and analyzed its dynamics during Mimivirus infection, along with the corresponding changes in the viral transcriptome. The AP transcriptome contains 13,043 genes, with N50 of 2,372 bp (Figure 2), and exhibits a median intron length of 72 bp and a canonical eukaryotic splice site sequence, for both donor and acceptor sequences [34] (Figure 3).

A summary of the key findings of the AP and Mimivirus functional enrichment analyses, as well as the state of the infected cells along the infection, is presented in Figure 7. Some Mimivirus genes were assigned to a different infection stage (early/intermediate/late) than those reported for Mimivirus infection of the related amoeba *Acanthamoeba castellanii* [12]. This may be due to differences between the host species or their multiplicity of infection (MOI), or differences in the sampling times.

**Figure 7.**
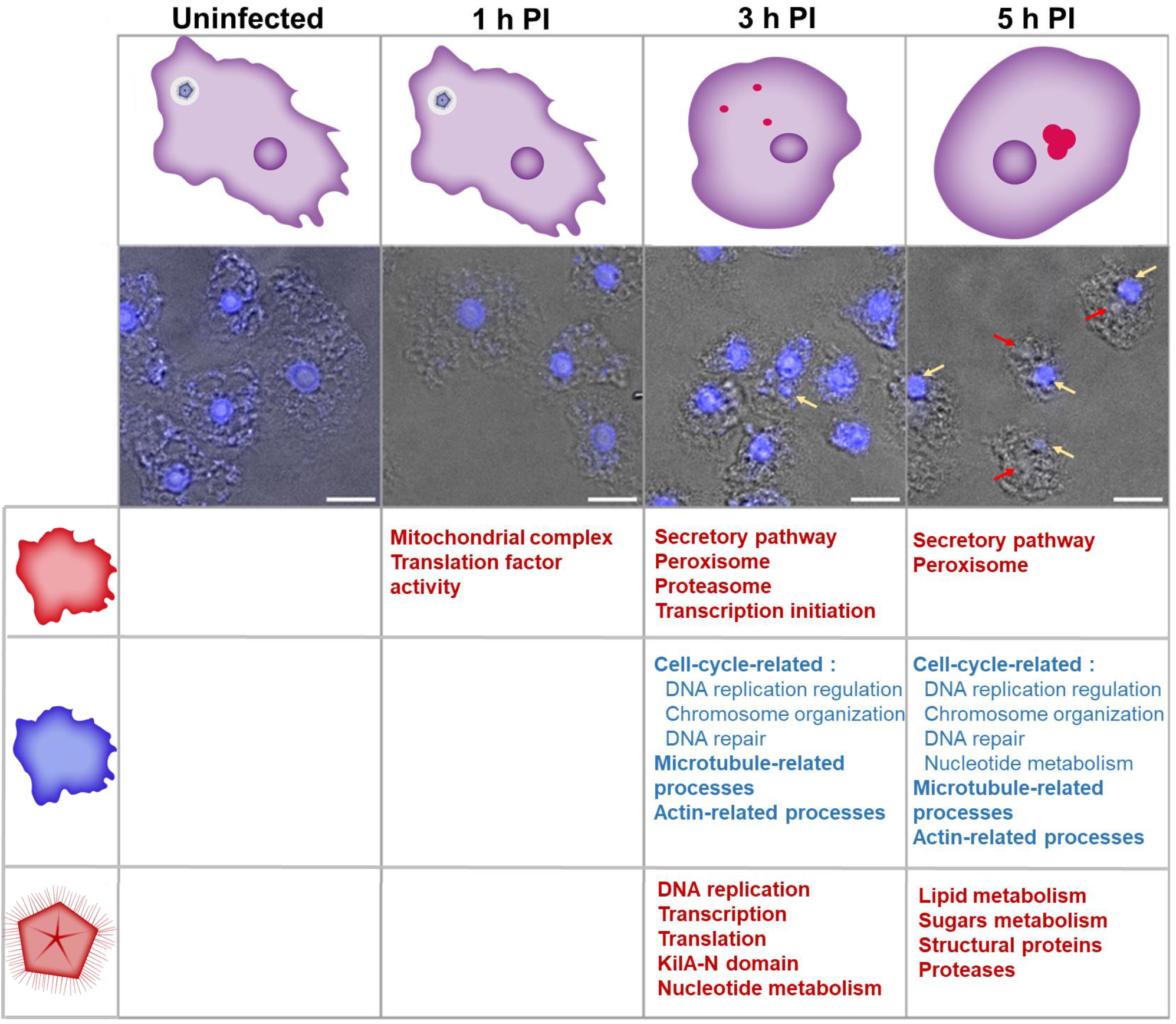
Summary of the *Acanthaomeba polyphaga* (AP) and *Acanthaomeba polyphaga mimivirus* (Mimivirus) transcriptome enrichment analysis. Each column represents a time point post-infection, as indicated on top. Upper panel: cell state illustration showing the nucleus (dark purple), viral factory (red), and Mimivirus (blue pentagon in a while circle). Lower panel: transmission light microscopy with 4′,6- diamidino-2-phenylindole (DAPI) labeling. Scale bar: 10 µm. Table at bottom: Summary of the processes that were upregulated in AP 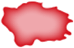, downregulated in AP 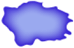, and upregulated in Mimivirus 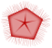.

Out of the 31 AP DEGs identified at 1 HPI, 29 were upregulated. These included genes associated with translation regulation and mitochondrial transport and morphogenesis (Figure 5B and C). Twenty-eight of the 31 genes were also differentially expressed and upregulated 1 h after UV stress (Supplementary Table 12). This suggests that the amoeba detects the virus early on during the infection and launches a general stress response. AP possesses genes that are annotated to immunity and pathogen recognition, such as NOD-like receptors (NLRs) LBP/BPI, leucine-rich repeat receptor-like protein kinase, TIR domain-containing protein, endoribonuclease dicer homolog 1, and toll/interleukin-1 receptor domain-containing protein [5], which may be involved in Mimivirus detection and the subsequent stress response.

Major changes in AP gene transcription patterns begin at 3 HPI (Figure 5B–D). During infection, the AP cells stop dividing (Supplementary Video 1). Based on our analysis, we suggest that the infection interferes with cell cycle progression by obstructing transition into the synthesis (S) and mitotic (M) phases. During the S phase, the cell is engaged in the execution of DNA replication [50, 51]. However, during infection, we observed a broad downregulation of host genes involved in DNA replication, DNA repair, and nucleotide metabolism. Our analysis similarly indicates enrichment in DRGs related to various mitotic processes, including chromosome organization and segregation, chromatin regulation and the formation of the anaphase-promoting complex (Figure 5A-D). By contrast, Mimivirus categories of DNA replication and nucleotide metabolism were enriched at 3 HPI (Figure 5A). In addition, we show that nucleotides used during replication of the viral genome originate primarily from *de novo* synthesis and not from breakdown of the host DNA or cellular reservoirs. Modulation of the cell cycle is by no means unique to the Mimivirus; infection by herpes simplex virus-1 (HSP1), Epstein-Barr virus [52], influenza A virus [52, 53], HIV, and SARS CoV [52] all lead to cell cycle arrest, typically at the G2\M phase. This may be an example of the advantage of the Mimivirus’s elaborate DNA replication system, which endows the virus with more autonomy in cases where genes associated with similar host tasks are downregulated.

Following a UV stress, there was also enrichment in AP DRGs associated with cell cycle-related categories that feature the GO terms cell cycle (GO:0007049), DNA replication (GO:0006260), mitotic sister chromatid segregation (GO:0000070) and anaphase-promoting complex (GO:0005680) (Supplementary Table 13). This suggests that cell cycle arrest is part of a general cellular stress-response of the host. Nonetheless, the arrest might be beneficial for the Mimivirus infection cycle, as it leaves metabolic resources, such as nucleotide precursors and amino acids, available for the virus.

The enrichment in DRGs related to the cytoskeleton at 3 and 5 HPI is consistent with microscopic observations of alterations in actin and tubulin structures during Mimivirus infection [11]. The rounding of cells and loss of the pseudopodia during infection are aligned with this notion and can be attributed to the downregulation of actin, myosin and tubulin-related processes (Figure 5B-D). However, actin is required for the proper fusion of VFs [11]. Thus, although actin-related processes are enriched in DRGs, actin is still required and is, therefore, likely present in amounts and forms that enable completion of the infection cycle.

It has been proposed that the Mimivirus membrane originates from the ER [13, 14]. We suggest that the mechanism by which Mimivirus induces the uptake of ER membranes relies on enhanced activation of the secretory pathway, as there is enrichment in UR genes related to that pathway. Viruses are known to utilize the ER for various tasks, such as entry into the proper cellular compartment, exiting the cell, viral replication and assembly, synthesis of glycoprotein [54, 55], and membrane assembly [13, 14, 54].

Our analysis also reveals enrichment in host UR genes associated with the ubiquitin-proteasome system, commencing at 3 HPI. The Mimivirus may utilize this system similarly to other viruses, such as HIV, HPV, influenza A virus and adenovirus, which use it to degrade host proteins, or cytomegalovirus (CMV), which exploits the ERAD pathway [56, 57]. Notably, the Mimivirus also possesses genes encoding proteins belonging to the ubiquitin-proteasome system (Supplementary Table 8), suggesting viral use of the system.

Finally, our analysis highlights changes in peroxisomal activities during infection. Other viruses were reported to modulate peroxisome numbers and activity upon infection [58, 59]. These include influenza A virus, which increases lipid metabolism in the peroxisome, and HCMV, HSV-1 and KSHV, which were associated with an increase in the number of peroxisomes in the host cells [58].

In conclusion, we propose that the host detects and responds to the Mimivirus early on during the infection; a result of this response may be cell cycle arrest observed during Mimivirus infection. In addition, we suggest that infection is accompanied by proteasome and peroxisome involvement in Mimivirus infection, given the increase in secretory pathway activity and the downregulation of actin and tubulin polymerization-related genes. The modulation in the latter probably underlines the massive alterations in the cytoskeleton of host cells during infection.

### Potential implications

Our work makes the *Acanthamoeba polyphaga* transcriptome publicly available. In addition, it provides leads for future research into the specific role of host cellular components, such as the peroxisome and ubiquitin-proteasome system, during Mimivirus infection. Future work should also focus on the sensing mechanisms used by the host to detect the Mimivirus at early stages of infection and on the molecular events that lead to cell cycle arrest of the host during infection, as indicated by our analysis.

## Methods

### Mimivirus purification

AP cells were plated in 175 cm^2^ flasks and infected by exposure to the virus for 48 h. The content of the flasks was then transferred to 50 mL tubes and centrifuged (100 g, 20 min, 4°C). The supernatant was transferred to a new 50 mL tube and centrifuged (10,000 g, 30 min, 4°C). The supernatant was discarded and virus pellets were washed twice with phosphate-buffered saline (PBS). The pellets were resuspended in PBS, and kanamycin (0.5 mg/mL) and ampicillin (1 mg/mL) were added. Viruses were stored at 4°C.

### Host cell growth and infection

AP cells were grown in peptone-yeast-glucose (PYG) in the presence of kanamycin (0.05 mg/mL) and ampicillin (0.1 mg/mL) in 9-cm untreated cell-culture plates to 70–80% confluency. AP cells were infected with Mimivirus using a multiplicity of infection (MOI) of 10. At 0.5 HPI, plates were washed with fresh medium and incubated for the desired infection time. Cells were then scraped from plates and frozen in liquid nitrogen until the RNA purification step. For the UV-stress experiment, cells were plated in a similar manner and exposed to 50 J/m^2^ UV light. After 1 h, cells were scraped and frozen.

### Illumina short read sequencing

RNA was purified from (1) uninfected AP samples, (2) AP samples at 1, 3, 5 HPI, and (3) AP samples 1 h after UV exposure, using RNeasy Mini Kit (Qiagen catalog number: 74104). RNA integrity was assessed using the 4200 TapeStation System. For library preparation, the poly-A fraction of mRNA was purified from 500 ng total RNA, followed by heat fragmentation and production of double-stranded cDNA. Thereafter, Agencourt Ampure XP bead cleanup (Beckman Coulter), end repair, A base addition, adapter ligation, and PCR amplification steps were performed. Libraries were quantified by Qubit (Thermo Fisher Scientific) and TapeStation (Agilent). Sequencing was performed on a Hiseq2500 instrument (Illumina) using a PE125 V4 kit (paired end sequencing).

### PacBio single-molecule real-time (SMRT) sequencing of long reads

RNA was purified from (1) uninfected AP samples, (2) AP samples at 1, 3, 5 HPI, and (3) AP samples 1 h after UV exposure, using RNeasy Mini Kit (Qiagen catalog number: 74104), and mixed. SMARTer® PCR cDNA synthesis kit (TaKaRa,cat 634926) and PrimeSTAR® GXL DNA polymerase (TaKaRa, cat R050B) were used for cDNA preparation. Libraries were prepared using the SMRTbell® Express Template Prep Kit 2.0. Sequencing was then performed using a SMRT cell 1M v3 on the Sequel instrument, using Sequel sequencing plate 3.0.

### Transcriptome assembly

Raw data obtained using PacBio were subjected to IsoSeq3 (version 8) for raw reads’ processing [16]. Illumina reads were mapped to PacBio reads using Bowtie2 [17, 18]. Unmapped reads were utilized using two independent approaches. The first was genome-based *de novo* assembly using StringTie software [19, 20] and the publicly-available genome assembly of AP (INSDC: CDFK CDFK00000000.1) [31]. We only used transcripts that originated from DNA contigs longer than 2000 nucleotides that were at least 10 nucleotides away from each end of the contig. The second approach involved *de novo* assembly without utilizing a reference genome, using rnaSPAdes [21, 22] (The names of transcripts originated from IsoSeq3, StringTie and rnaSPAdes begin with “UnnamedSample_HQ_transcript_”, “new.strg.” and “NODE_”, respectively). Transcripts from the three assemblies were integrated and EvidentialGene, tr2aacds [23, 24] was used to remove redundancy. The transcriptome was run against the non-redundant protein database (NCBI) using Blastx [25], for 20 hits, and the best hit results were used for further analysis. The NCBI Taxonomy browser [60] was used to analyze the organisms from which the annotation was borrowed, based on the Blastx results. In order to remove transcripts originating from contaminants, we filtered Blastx results for bacteria and removed those that had a hit of >85% identity and a coverage > 50%. Transcripts that aligned with viruses were likewise filtered out. The Cogent - collapse_by_cogent_category.py script [27] was later used to define genes, and the longest transcript was chosen as the family representative.

### Splice junction analysis

Transcripts generated by IsoSeq3 were mapped to AP genome contigs of length >10 kb, using the minimap2 software [32, 33]. Unmapped reads were removed from the same files using samtools view – F4. Tama scripts (https://bmcgenomics.biomedcentral.com/articles/10.1186/s12864-020-07123-7) were applied to create a gtf file from the alignment (tama_collapse.py -i 95 and tama_convert_bed_gtf_ensembl_no_cds.py). Only in cases where a full transcript was mapped to the genome contig were the splice junction coordinates extracted from the gtf file. Redundancy was removed and the sequences were extracted with bedtools getfasta[57, 61]. A nucleotide representation logo was created using WebLogo [62].

### Quality assessment of the assembly

Mimivirus transcriptome assembly was used in order to assess the quality of the transcriptome assembly. Transcripts originating from the Mimivirus were aligned to the published Mimivirus transcriptome using BLAT [63, 64] and the percentage of covered and duplicated transcripts was calculated. In addition, BUSCO [36, 37] was used to evaluate the completeness and duplication of the transcriptome. In order to assess the quality of the assembly, we compared our *de novo* AP transcriptome BUSCO results with those of its closest relative with a known transcriptome, namely, AC. We used both eukaryota_odb9 and protists_ensembl databases for the comparison.

### Sanger sequencing

The KAPA Taq PCR Kit (Sigma-Aldrich KR0352_S) was used to amplify the reads from a cDNA mixture of uninfected samples and 1, 3 and 5 HPI samples. Primers were designed using the Primer3 software [65, 66] (Supplementary Table 14). The forward primers were used in the sequencing (3730 DNA Analyzer, ABI). DEGs with Blastx annotations from each assembly tool were selected randomly.

### Mapping and gene quantification

The RSEM tool [67] was used to map Illumina reads from each sample to a unified transcriptome that included the Mimivirus and AP (with isoforms). Differential expression analysis was applied at the gene level by summing the number of reads assigned to all isoforms of the same gene. The fold-change value and the p-value between each two time points were calculated using the DESeq2 package in R [68]. The criteria for DEGs were ≥ 2-fold change and a false discovery rate (FDR) (adjusted p-value) <0.05 in at least one comparison between the samples.

### Data clustering and PCA

DESeq2 output of normalized rlog transformed (rld) values were used for principal component analysis (PCA), performed with Partek Genomics Suite software (version 6.5),(Partek Inc., St. Louis, MO). K-means clustering was performed using standardized log2 DEG expression values as input (rld). A Silhouette plot was used to determine the optimal number of clusters for the Mimivirus data (Matlab) (Supplementary Figure 2), and clustering was then performed using the Partek Genomics Suite.

### Gene annotation by GO

The Blast2Go tool [29] was used to find GO-terms associated with the hit sequences obtained by Blastx (against the non-redundant protein database [69]). The output was analyzed by Ontologizer (Supplementary File 1).

### Biological pathways analysis

GO functional enrichment analysis for the annotated genes in the different clusters was performed using Ontologizer [44, 45] and the Obo library [70]. The topology weighted algorithm [71] was used and the threshold p-value was 0.005. The assignment of genes to GO terms can be found in Supplementary File 4. GO terms in each domain were grouped into categories in a semi-automatic manner, using a combination of Revigo [72] and manual classification, in order to simplify data interpretation. Manual classification was based on GO hierarchical classification and common genes between GO terms. The complete list of enriched GO terms and the categories into which they were grouped are provided in Supplementary Table 8.

### Mimivirus enrichment analysis

Mimivirus genes were categorized manually, based on the top blast annotations and the literature (Supplementary Table 1). Category counts were performed using an in-house script (doi 10.6084/m9.figshare.20154740). The enrichment analysis was performed using hypergeometric test, in R. Enrichment was considered statistically significant with a p-value < 0.05.

### Fluorescence microscopy

Uninfected AP cells were plated on 3.5-cm glass-bottomed plates (µ-Slide, 8-well, tissue culture-treated (Ibidi IBD-80826 i)). Cells were infected with Mimivirus at MOI∼10 for 1, 3 or 5 h. The plates were then washed and incubated with PFA 4% in PBS for 20 min at room temperature (RT), washed three times with PBS for 5 min each, and incubated for an additional 10 min at RT with triton X100 0.1% in PBS. Cells were washed three times with PBS for 5 min each and incubated with a 1:1000 dilution of DAPI (5 µg/mL) in PBS for ∼1 h, and then finally washed three times with PBS. Samples were imaged using an Eclipse TI-E Nikon inverted microscope (Nikon Instruments Inc., Melville, NY) with a Plan Apo 100X/1.4 NA lens. Images were acquired with a cooled electron-multiplying charge-coupled device (EMCCD) camera (IXON ULTRA 888; Andor).

### Amoebal genomic DNA staining by EdU

We used the Click-iT® EdU Cell Proliferation Kit for Imaging (Thermo Fischer, catalog number: C10337). AP cells (5–10% confluency) were grown in a 3.5-cm petri dish containing EdU nucleotides (20 μM; 1:500), for 24 h at 30°C. Cells were transferred to 8-well plates (Ibidi 80826), to achieve 50–70% confluency, for 10 minutes, and then washed thrice with fresh PYG and left in the incubator for 1 h, followed by Mimivirus infection at MOI = 10 for 4, 5, 6, 8, and 10 h. Infection was terminated by washing the cells with PBS and incubating them in formaldehyde (300 μL, 3.7% in PBS) for 15 minutes, followed by three washes with PBS. Triton® X-100 (300 μL, 0.1% in PBS) was added, and the cells were then incubated for an additional 10 minutes at RT, after which they were washed thrice with PBS. We then followed the manufacturer’s instructions for using the kit. Briefly, 300 μL of Click-iT® reaction mixture was added to each well and the cells were incubated for 30 minutes at RT, protected from light. The reaction mixture was then removed and the cells were washed three times with 3% BSA in PBS and stained with DAPI 1:1000.

### Time-lapse imaging during Mimivirus infection

AP cells were grown on 35 mm^2^ glass-bottom plates (Mat-Tek corp. P35G-1.5-14-C), treated with 1 mg/mL poly-L lysine for 1 h, and then infected with Mimivirus at MOI ∼50, at 30°C. Widefield images were acquired using a Deltavision microscope (Applied Precision) equipped with a 60X UPlanSApo NA 1.40 objective and a photometrics coolSNAP HQ2 CCD (Roper Scientific, Tucson, AZ). Images were captured every 60 s from 250 min post-infection onward.

### AP protein database

The database (Supplementary File 3) was created on the basis of the *de novo* AP transcriptome. The transcripts were translated using in-house Perl scripts. Each transcript was translated in six frames to determine potential ORFs, using the EMBOSS function transeq (http://emboss.sourceforge.net/apps/release/6.0/emboss/apps/transeq.html). A BlastX search against a non-redundant protein database was performed for each transcript and the first three BlastX hits were used to determine the frame and coordinates of the ORF. If two or three of the obtained blast hits were of the same reading frame, the frame and coordinates were compared to the putative ORFs and selected. When reading frames were different in each BlastX hit, ORFs of these frames were checked and the longest ORF was selected. If there were no BlastX hits, the longest ORF was chosen. Only proteins containing more than 49 aa were included in the database.

## Availability of Supporting Data and Materials

The data sets supporting the results of this article are available in the Figshare repository, 10.6084/m9.figshare.20154740.

Raw data have been deposited in the NCBI BioProject database, BioProject accession number PRJNA720295 (https://dataview.ncbi.nlm.nih.gov/object/PRJNA720295).

## Abbreviations

AC: Acanthamoeba castellanii
AP: Acanthamoeba polyphaga
bp: base pairs
BUSCO: Benchmarking Universal Single-Copy Ortholog
DEGs: differentially expressed genes
DR: downregulated
ER: endoplasmic reticulum
ERAD: ER- associated degradation
FDR: false discovery rate
GO: Gene Ontology
HIV: human immunodeficiency virus
HPV: human papilloma virus
CMV: cytomegalovirus
LR: long read
NCLDV: nucleo-cytoplasmatic large DNA virus
NGS: next-generation sequencing
NLR: NOD-like receptors
ORF: open reading frame
PCA: principal component analysis
HPI: hours post infection
RT: room temperature
SMRT: single-molecule, real-time
SR: short reads
TF: transcription factor
URG: upregulated genes
VF: viral factory
VV: vaccinia virus

## Authors’ contributions

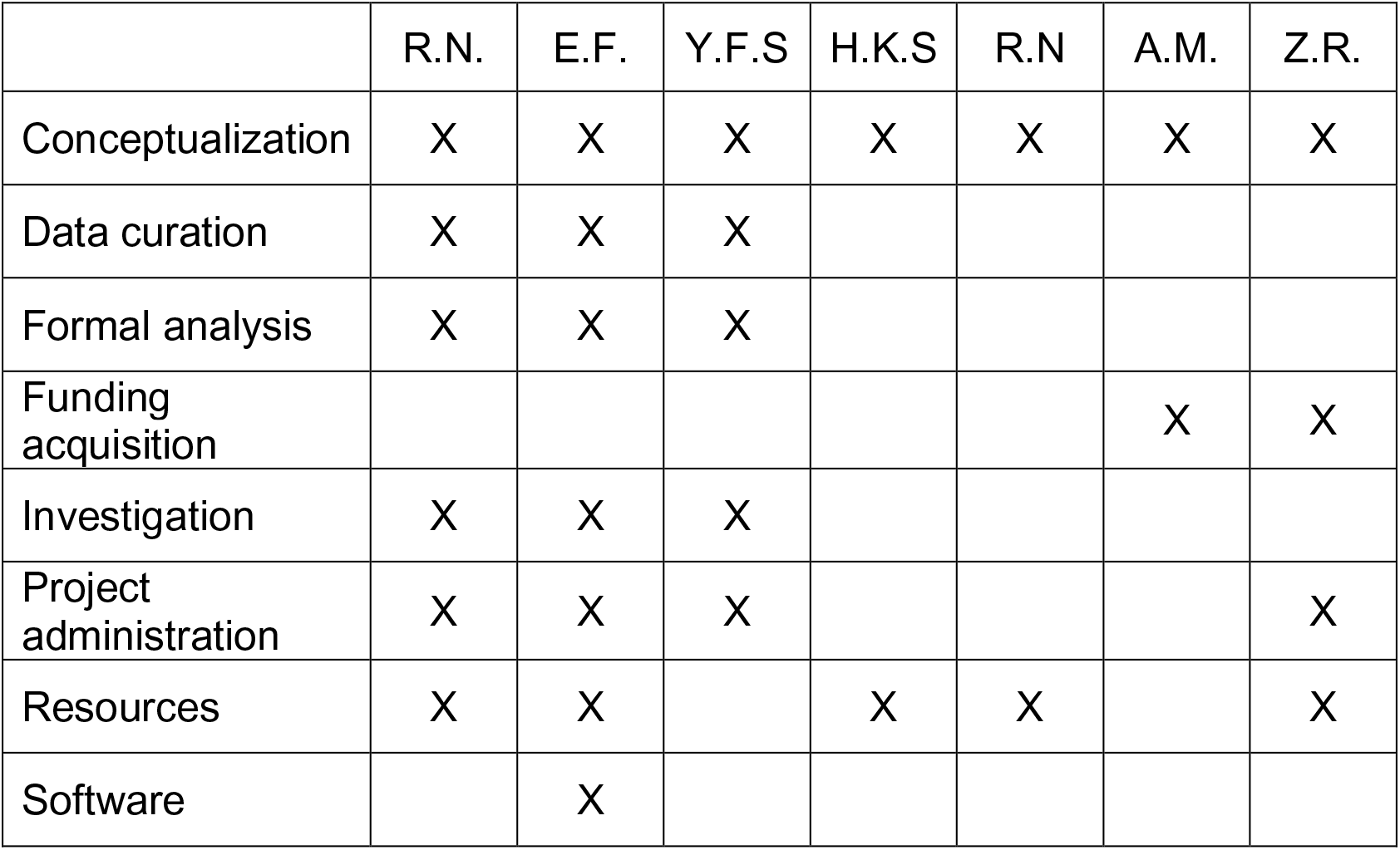
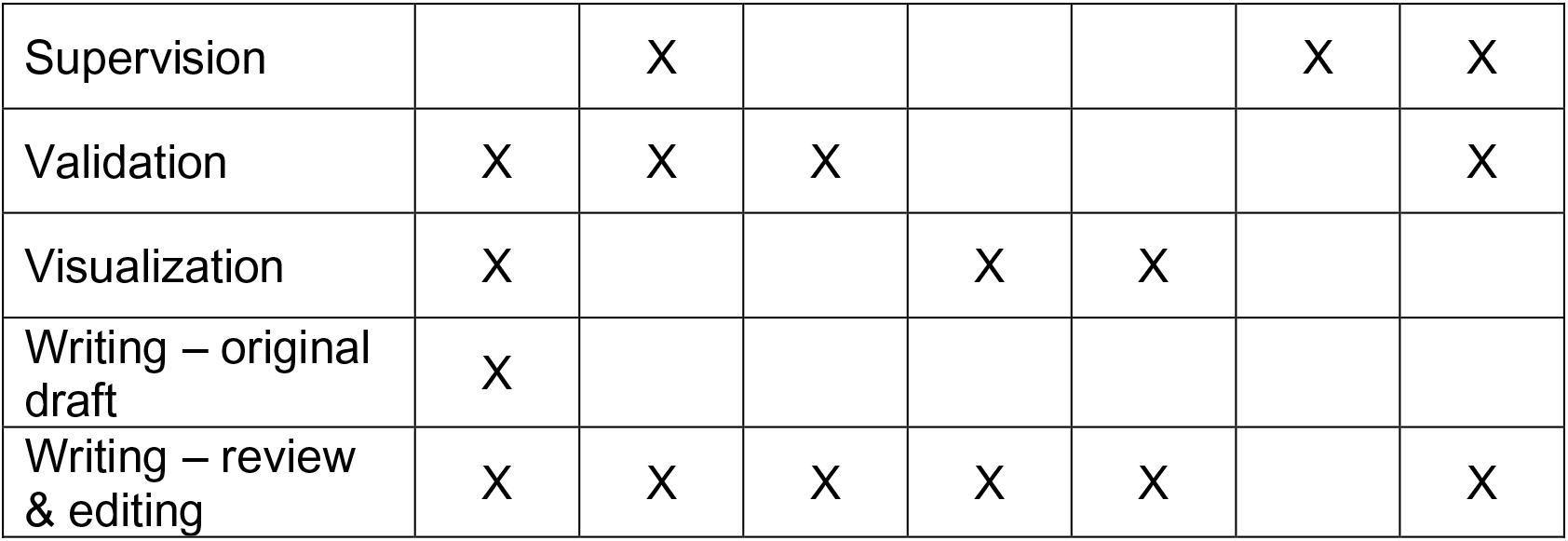

## Acknowledgements

We thank Drs. Merav Kedmi and David Pilzer, for their support in SMRT sequencing sample preparation, Maor Knafo, for his assistance with the SMRT sequencing and data analysis, and Drs. Ruti Kapon and Inbal Neta-Sharir, for their helpful advice and for reviewing the manuscript.

## Supplementary material

10.6084/m9.figshare.20154740

